# Analysis of somatic mutations in senescent cells using single-cell whole-genome sequencing

**DOI:** 10.1101/2022.09.16.508266

**Authors:** Lei Zhang, Marco De Cecco, Moonsook Lee, Xiaoxiao Hao, Alexander Y. Maslov, Cristina Montagna, Judith Campisi, Xiao Dong, John M. Sedivy, Jan Vijg

## Abstract

Somatic mutations accumulate in multiple organs and tissues during aging and are a known cause of cancer. Here we tested whether mutations accumulate during replicative senescence. Cellular senescence is also a possible cause of functional decline in aging, yet also acts as an anti-cancer mechanism *in vivo*. Using single-cell whole-genome sequencing, we compared mutation burdens between early passage and deeply senescent human fibroblasts. The results showed that single-nucleotide variations and small insertions and deletions increased in senescent cells by about two-fold, but have the same spectrum as early passage cells, while it has been known that particular mutational signatures are found in tumor cells. In contrast, aneuploidies were observed in half the senescent cells, but largely absent in early passage cells. Thus, the patterns of mutations among senescent, normal-aged and tumor cells differ significantly.

## Introduction

Life-long accumulation of mutations in somatic cells has been implicated as a driver of cancer and aging soon after the discovery of the structure of DNA (Failla, 1958; Szilard, 1959). Somatic mutations have been detected in clonal cell lineages, such as in cancer, but in the absence of a selective advantage, their random nature and low abundance has stymied accurate detection during aging. With the advent of single-cell whole-genome DNA sequencing, accumulation of mutations can be accurately documented and quantified in multiple tissues and cell types during human aging with or without clinical conditions (Vijg & Dong, 2020).

Cellular senescence is a cell fate elicited by a variety of stresses and characterized by an essentially irreversible arrest of proliferation (Gorgoulis et al., 2019). Irreparable DNA damage caused by telomere shortening or replication fork collapse due to oncogene activation are well-studied triggers, and in many of these contexts senescence has a beneficial role as a tumor suppression mechanism. Senescent cells however also accumulate sporadically in most tissues with aging, where they exert a strong proinflammatory influence (Coppé et al., 2008) and are believed to promote many age-related pathologies and diseases (Pignolo, Passos, Khosla, Tchkonia, & Kirkland, 2020). Senescence is a complex state that includes widespread gene expression and epigenetic changes (Cruickshanks et al., 2013), mitochondrial dysfunction, elevated reactive oxygen species (ROS) (Correia-Melo et al., 2016), and persistent DNA double-strand breaks (DSBs) and DNA damage response (DDR) signaling (Fumagalli et al., 2012).

Little is known about the pattern and role of mutations during cellular senescence. On one hand, given the evidence of ROS and DSBs, one might expect accelerated mutation rates in senescent cells; on the other hand, it is reasonable to hypothesize limited mutagenesis given that senescence is a major anti-cancer mechanism that prevents subsequent cell proliferation (Campisi, 2013). To test these conflicting hypotheses, we performed single-cell whole-genome sequencing of early passage and senescent normal human lung fibroblasts and compared the frequencies and spectra of three mutagenic processes: single-nucleotide variations (SNVs), small insertions and deletions (INDELs), and aneuploidies.

## Results

### Single-cell whole-genome sequencing of deeply senescent cells and controls

We generated replicatively senescent cells by continuous passaging of the IMR-90 normal human fibroblast cell strain. Since L1 retrotransposons are transcriptionally upregulated 3-4 months after this proliferative arrest (De Cecco et al., 2019), and to allow ample time for other mutagenic events to accumulate, we maintained the senescent cultures for five months (hereafter referred to as deep senescence). Single cells were isolated from both early passage and deeply senescent cells using the CellRaft System (Cell Microsystems) and amplified using the SCMDA procedure that we developed previously (Dong et al., 2017). We performed whole-genome sequencing (average depth of 24x; Table S1) of early passage and deeply senescent single-cell amplicons. We also sequenced bulk DNA from early passage cells, which we used to filter out germline polymorphisms (Figure 1). Somatic SNVs, INDELs and aneuploidies were identified from each single cell using the SCcaller and SCCNV pipelines that we developed previously (Dong, Zhang, Hao, Wang, & Vijg, 2020; Dong et al., 2017). Frequencies of SNVs and INDELs per cell were estimated after correcting for genome coverage and sensitivity in variant calling (Table S2) (Zhang et al., 2019).

**Figure 1.**
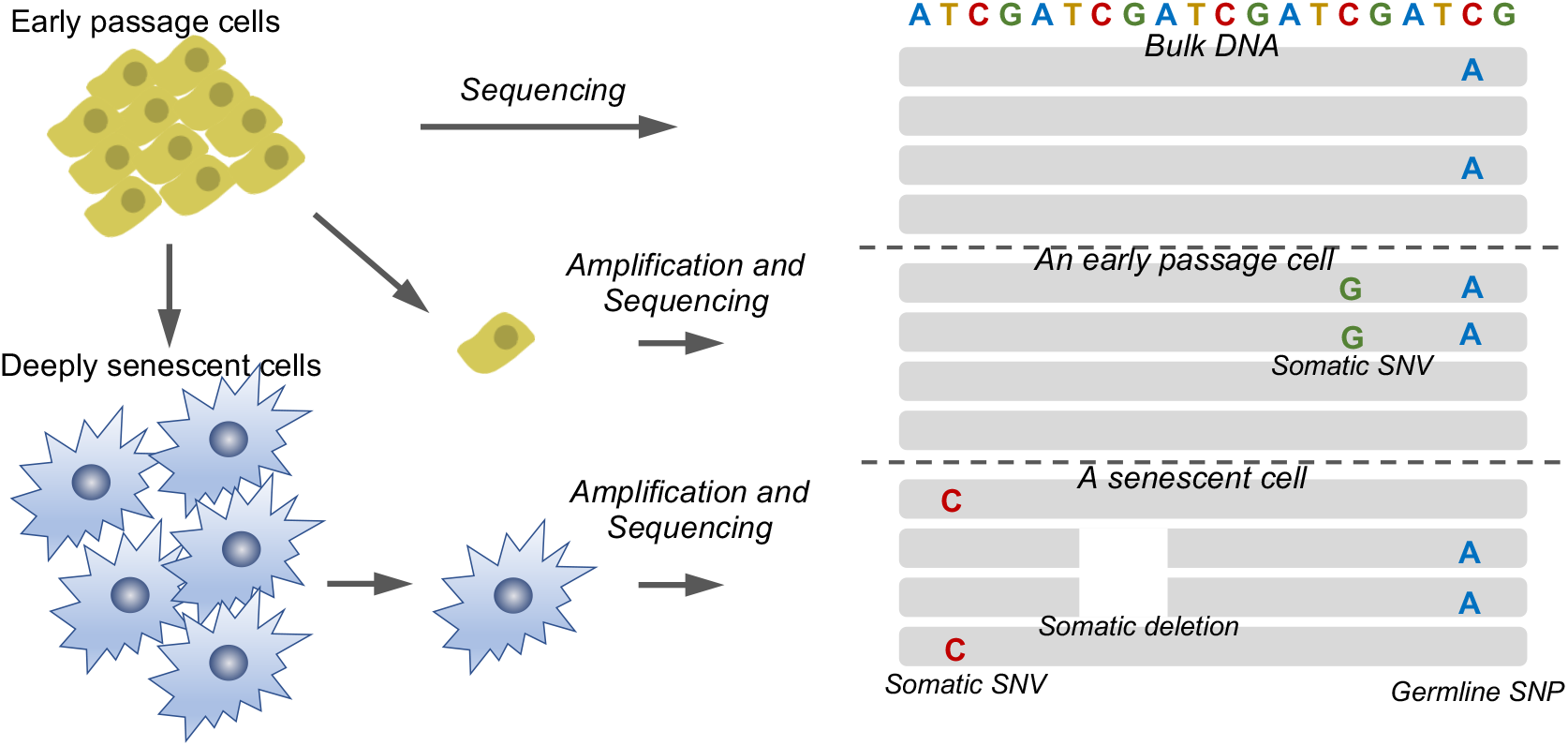
Schematic illustration of the detection of somatic mutations by single-cell whole genome sequencing. Each track in grey presents a sequence read, and a dotted line separates reads of different samples.

### Elevated frequencies of SNVs and INDELs in senescent cells

We observed a median of 1,103 SNVs per cell in early passage and 2,618 SNVs per cell in deeply senescent cells (*P*=0.0022, Wilcoxon Rank Sum Test, two-tailed; Figure 2A), reflecting a 2.4-fold higher SNV frequency in senescence. Of note, the frequency of 2,618 SNVs per cell is somewhat less but close to the 3,127 SNVs per cell we reported in primary B lymphocytes in human centenarians (Zhang et al., 2019). For INDELs we also observed significantly higher frequencies in senescent compared to early passage cells (median frequencies of 311 and 165 per cell type, respectively; 1.9-fold; *P*=0.0128, Wilcoxon Rank Sum Test, two-tailed; Figure 2B).

**Figure 2.**
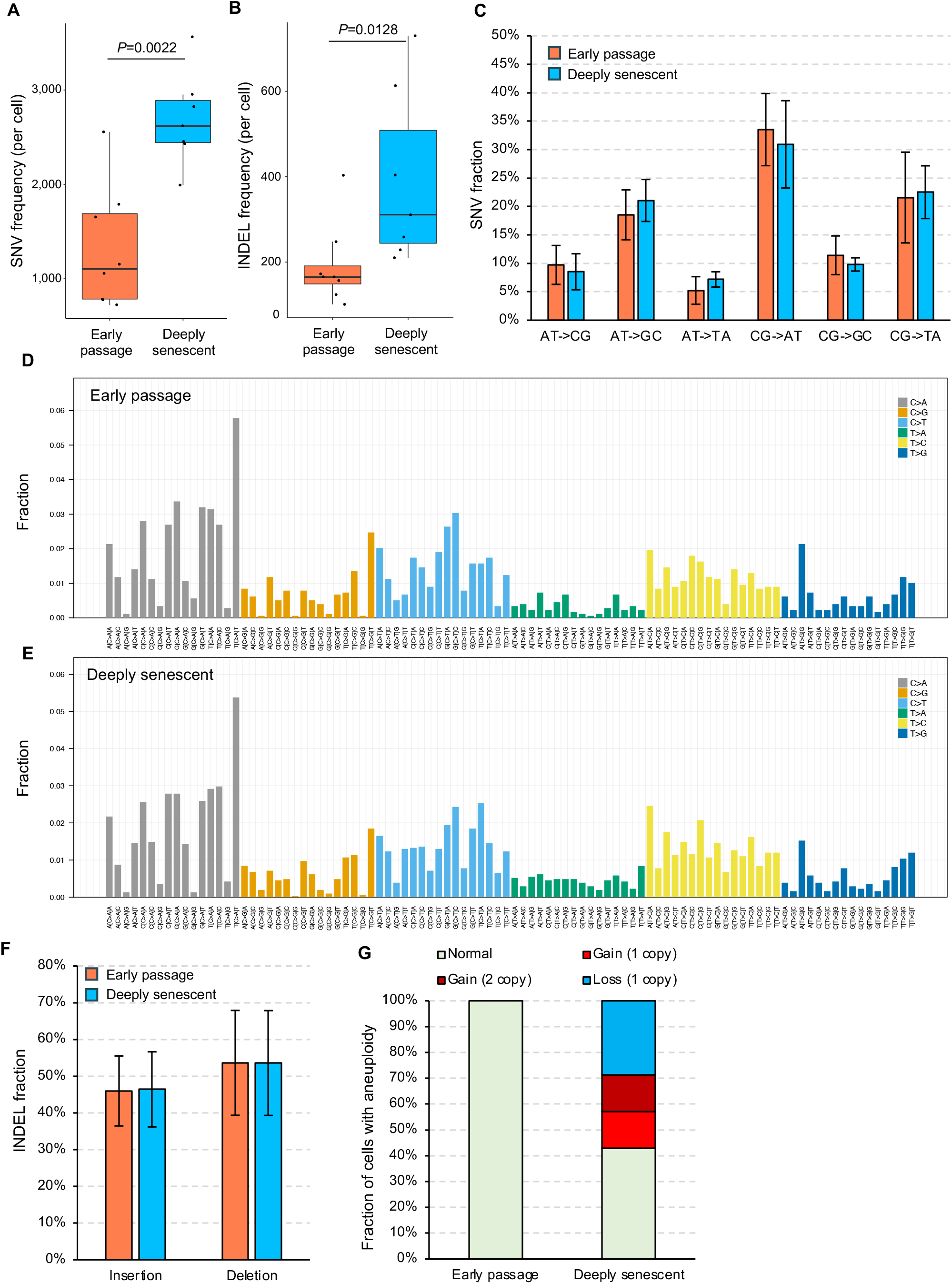
Somatic mutations in early passage and deeply senescent cells. SNV (**A**) and INDEL frequencies (**B**). SNV spectra (**C**). Refined SNV spectra by considering their flanking bases in early passage (**D**) and senescent cells (**E**) separately. INDEL spectra (**F**). Types of whole-chromosome aneuploidies (**G**). For (**A**) and (**B**), *P* values were estimated using the Wilcox Rank Sum test, two-tailed. Box plot elements are: center line, median; box limits, upper and lower quartiles; whiskers, 1.5× interquartile range; points, all data points. For (**C, F**), bars represent means and error bars standard deviations.

A higher load of SNVs and INDELs in senescent cells is expected since these types of mutations are typically a result of replication errors. To test if additional mutational mechanisms occur during senescence, we first compared mutation spectra of early passage and senescent cells. For SNVs, we observed no significant differences in the fraction of all six SNV types between the two conditions (Figure 2C). The highest fraction of mutations was CG>AT transversions, followed by CG>TA and AT>GC transitions. This similarity between early passage and senescent cells remained when considering the bases flanking the mutations (Figures 2D-E). For INDELs, we also observed no significant differences: deletions were 54% of all INDELs, with the remainder being insertions under both conditions (Figure 2F). In contrast, particular SNV and/or INDEL signatures are often found tumor cells (Alexandrov et al., 2020).

We next analyzed the genomic distribution of SNVs and INDELs. Plotting the density of mutations in early passage and senescent cells across the genome to identify possible mutations hotspots using the R package “karyoploteR” (Gel & Serra, 2017) did not reveal any significant chromosomal position enrichment of mutations (Supplementary Figures 1A-B), which is often found in tumor cells (Wang et al., 2007). This finding was confirmed using the mutation showers algorithm with the R package “ClusteredMutations” (Lora, 2016; Wang et al., 2007), which found no obvious mutational hotspots. Overall, the high similarities in SNV and INDEL spectra and chromosomal distributions between early passage and senescent cells indicates the absence of additional mechanisms for generating SNVs and INDELs in senescent cells.

In cancer as well as aging of the hematopoietic system, clonal development of somatic cell lineages was often observed and associated with health status of human subjects (Greaves & Maley, 2012; Jaiswal & Ebert, 2019). To test if clonal development can occur during passaging to replicative senescence, we compared the frequencies of overlapping mutations found in two or more cells. Because the mutation frequencies in these cells are not very high, an overlapping mutation found in two cells is unlikely to arise by two independent events, and much more likely the result of a mutation occurring in a progenitor cell. We found 4.6% of mutations in early passage cells occurred in more than one cell, and 10.8% in senescent cells (*P*<2.2×10^−16^, Chi-squared test, two tailed). This 2.4-fold increase suggests an increase in clonal development of mutations as cells approach senescence.

We also annotated the predicted effects of mutations in protein-coding sequences using wANNOVAR (Chang & Wang, 2012). Among the 65 non-synonymous or loss-of-function mutations observed, a stop codon mutation in the *SLC36A9* gene and non-synonymous mutation in the *CCDC114* gene are notable (Table S3). Both mutations were observed in three out of the seven senescent cells analyzed, indicating that they have expanded to a significant fraction of the population prior to senescence.

### Aneuploidies were only observed in senescent cells

We also able identified aneuploidies in the single-cell sequencing data. In early passage cells we did not observe any whole-chromosome aneuploidy, although we found one large deletion on one copy of chromosome 2 in one of the 8 cells (Supplementary Figure 2). In contrast, we observed that 4 of the 7 senescent cells have whole-chromosome gain or loss aneuploid, including a 2-copy-gain of chr. 18, a 1-copy-loss of chr. 19, a 1-copy-gain and 1-copy-loss of chr. 22 (Figure 2G and Supplementary Figure 3). These results indicate that unlike smaller mutations (SNVs and INDELs), the rate of aneuploidies is significantly elevated in senescent cells than early passage cells. As senescent cells permanently exit cell cycle, these excessive errors are likely due to errors in the last mitotic replication before the cells become senescent. This finding is in agreement with previous findings of increased aneuploidy in human senescent cells (Andriani et al., 2016) and suggests that aneuploidy accompanies the entry into senescence and can induce senescence (Andriani et al., 2016).

## Discussion

The role of somatic mutations as a cause of chronic conditions or diseases, such as cancer and aging, has been hypothesized since the 1950s. Soon afterwards, mutations were implicated as an important cause of cancer (Nowell, 1976). In normal somatic cells during aging, mutations were much more difficult to discover than those in tumor cells. This difficulty resides in the fact that most normal cells do not clonally expand, at least not to the extent of tumor cells. With the development of single-cell or clone sequencing, mutation accumulation has been discovered during human aging in all cell types analyzed (Blokzijl et al., 2016; Brazhnik et al., 2020; Lodato et al., 2018; Zhang et al., 2019). Mutation frequencies differ significantly by cell type, but often in the range of a few hundred or less per cell in newborns to a few thousand per cell in the elderly. The effect of these mutations varies by their specific loci: although some can provide a growth advantage to cells, e.g., in clonal hematopoiesis (Zink et al., 2017), most functional mutations were observed to be negatively selected during aging, at least in lymphocytes (Zhang et al., 2019).

In this study, we directly analyzed mutation frequencies and spectra in replicatively senescent human cells. The patterns of mutations differ substantially from tumor mutations in several aspects, suggesting different roles of mutations between tumor and senescent cells.

Senescent cells do not acquire SNVs and INDELs any more than early passage cells. SNVs and INDELs in senescent cells do not cluster at potential hotspots and their increase in mutation frequencies are in the same range as those in normal cells during aging, suggesting the absence of significant driver mutations, which are typically found in tumors. Still, SNVs and INDELs can occur at locations potentially damaging to cellular function, e.g., the mutations observed at *SLC36A9* and *CCDC114*. Germline mutations in both genes were known to be associated with damage in lung (Corvol et al., 2018; Onoufriadis et al., 2013), which is the tissue type that the cell line was isolated from. In addition, the excessive number of aneuploidies (more than 50% of cells) observed in senescent cells was substantially higher than normal cells during aging, which are often loss of sex chromosomes (Forsberg, 2017; Machiela et al., 2016), but high frequency of aneuploidy are observed in most tumors (Ben-David & Amon, 2020).

Overall, the above results provide the first evidence of somatic mutation accumulation in senescent cells. They suggest that the effect of mutations in senescent cells is less damaging than those in tumor cells, but more than those in normal cells in aging. As mentioned above, senescence is a comprehensive phenotype and can be a result of many different factors besides replication. It still requires future investigation whether the patterns and roles of mutations are the same or different in different types of senescent cells.

## Experimental Procedures

IMR-90 fibroblasts were obtained from the ATCC (CCL-186) and cultured using physiological oxygen conditions (92.5% N_2_, 5% CO_2_, 2.5% O_2_) as described (De Cecco et al., 2019). Cultures were serially propagated at 1:4 dilution at each passage (hence 1 passage is equivalent to 2 population doublings). After proliferation ceased (passage 38) the cultures were kept in the senescent state for 4 months with regular media changes as described (De Cecco et al., 2019).

Single cells were isolated into individual PCR tubes using the CellRaft System (Cell Microsystems) according to manufacturer’s instructions and kept at -80 °C until amplification.

Single-cell whole-genome amplifications were performed by using the single-cell multiple displacement amplification (SCMDA) method (Dong et al., 2017). In brief, 1 μl Exo-Resistant random primer (Thermo Fisher Scientific) and 3 μl lysis buffer (400 mM KOH, 100 mM DTT and 10 mM EDTA) were added to the cell and kept on ice for 10 min. Then, 3 μl stop buffer (400 mM HCI and 600 mM Tris-HCI pH 7.5) was added for neutralization. Finally, 32 μl Master Mix containing 30 μl MDA reaction buffer and 2 μl Phi29 polymerase (REPLI-g UltraFast Mini Kit; Qiagen) were added. The amplification was performed at 30 °C for 1.5 hr, 65 °C for 3 min, and 4 °C until purification. Purification was performed using AMPureXP-beads (Beckman Coulter). The final product was quantified using the Qubit High-Sensitivity dsDNA kit (Thermo Fisher Scientific). Bulk DNA was extracted from early passage fibroblasts using the Quick-DNA kit (Zymo Research).

Whole-genome sequencing libraries were prepared using the TruSeq Nano DNA HT Sample Prep Kit (Illumina), and sequenced on the Illumina HiSeq X Ten System with 2×150 bp paired-end reads by Novogene. After quality control and trimming using FastQC (v0.11.9) and Trim Galore (v0.6.4) (Gossen et al., 1989; Kong et al., 2012), sequencing reads were aligned to the human reference (Build 37) using BWA mem (v0.7.17) (Li, 2013). PCR duplicates were removed using samtools (v1.9) (Li et al., 2009). The alignment was INDEL-realigned and base pair recalibrated using GATK (v3.5.0) (McKenna et al., 2010). Somatic SNVs and INDELs of a single cell were called using SCcaller (v2.0.0) requiring at least 20x sequencing depth and by comparing the single-cell data to the bulk DNA-seq data (Dong et al., 2017). Somatic RTs were called using TraFiC-mem with default parameters (Rodriguez-Martin et al., 2020). Large copy number variations and aneuploidies were identified using SCCNV (bin size: 500k bp; no. bins per window: 100; v1.0.2) (Dong et al., 2020).

## Acknowledgements

This work was supported by the American Federation For Aging Research Grant (the Sagol Network GerOmic Award for Junior Faculty for L.Z.), the NIH NIA R00 AG056656 (X.D.), R01 AG016694 (J.M.S.), P01 AG051449 (J.M.S.), P01 AG017242 (J.V.), P01 AG047200 (J.V.), and U01 ES029519 (J.V.).

## Author contributions

J.M.S. and J.V. conceived the study, L.Z., M.D.C. and M.L. performed experiments, X.D. and X.H. analyzed the data, and L.Z. and X.D. prepared the manuscript with input from all co-authors.

## Data availability statement

Single-cell sequencing data will be submitted to NCBI dbGAP database before publication.

## Supporting Information Listing

**Supplementary Figure 1. Rainfall plot illustrating distances of neighboring mutations**. Mutations of single cells from each group were pooled in this analysis. (Top) Density of mutations in kilobase bins. Distances of each mutation to its closest other mutation.

**Supplementary Figure 2. Aneuploidy and large copy number variation in early passage single cells**.

Each panel indicates one single cell. Curved lines indicate normalized sequencing depth, and horizontal straight lines indicate estimated copy numbers. Of note, some parts of the genome do not have an estimated copy number. This is due to high fluctuation of sequencing depth in these regions, and the SCCNV pipeline did not report their copy numbers due to lack of statistical confidence. This is a typical problem of single-cell whole-genome amplification, and affects mainly copy number estimation. The arrow indicates a large copy deletion in chromosome 2 of a single cell.

**Supplementary Figure 3. Aneuploidy and large copy number variation in deep senescent single cells**.

The four black arrows indicate four aneuploidies observed in the single cells. In addition to the four events, cell “IMR90_P38_02_10” has potentially a copy-number-loss on chromosome 22 (blue arrow), although it is not called by SCCNV due to the high fluctuation in sequencing depth in this single cell. The fluctuation of sequencing depth is a result of whole-genome amplification, which affects mainly copy number estimation.

