## Supplemental Figures 1-3 for "Analysis of somatic mutations in senescent cells using single-cell whole-genome sequencing"

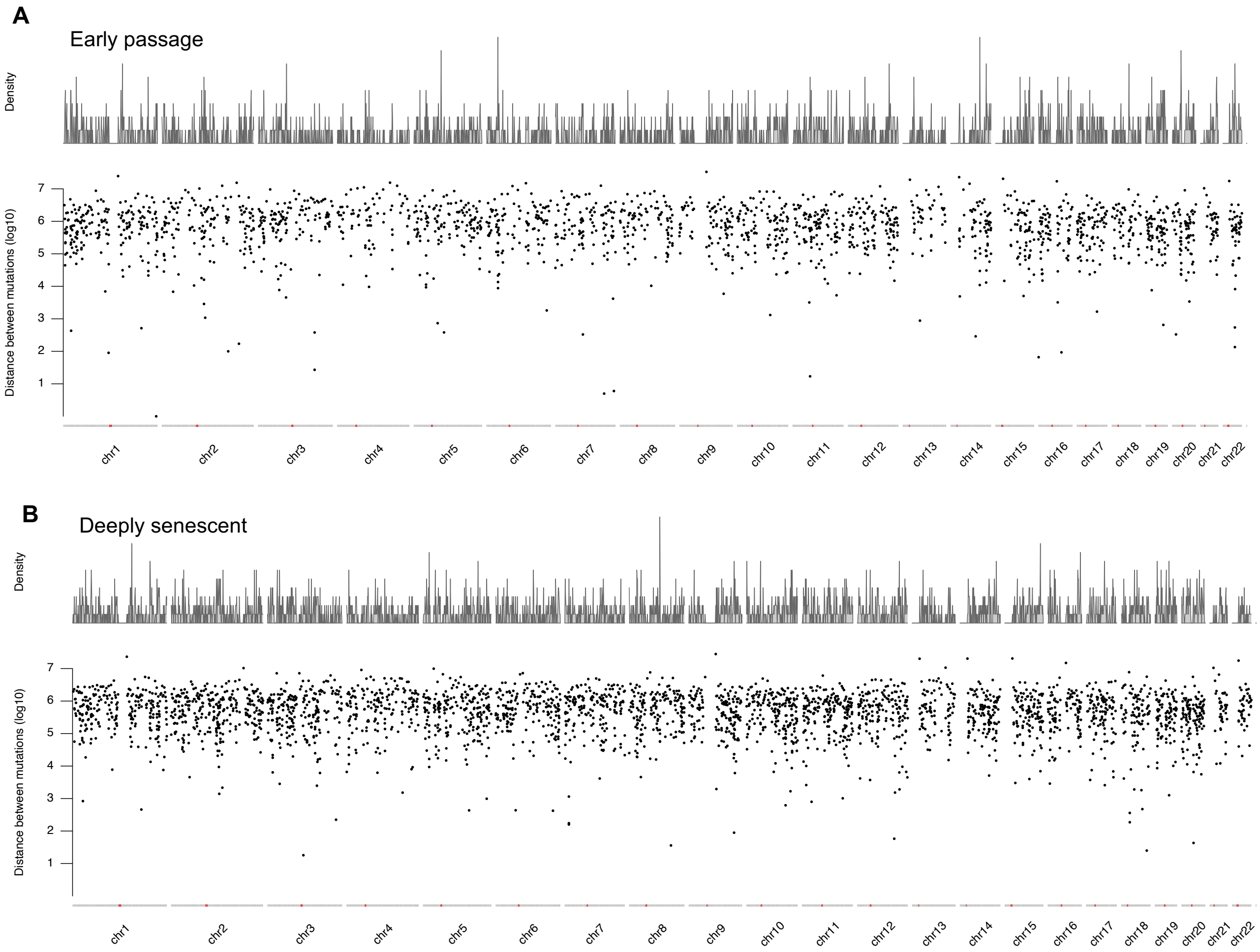


**Supplementary Figure 1.**

Early passage cells

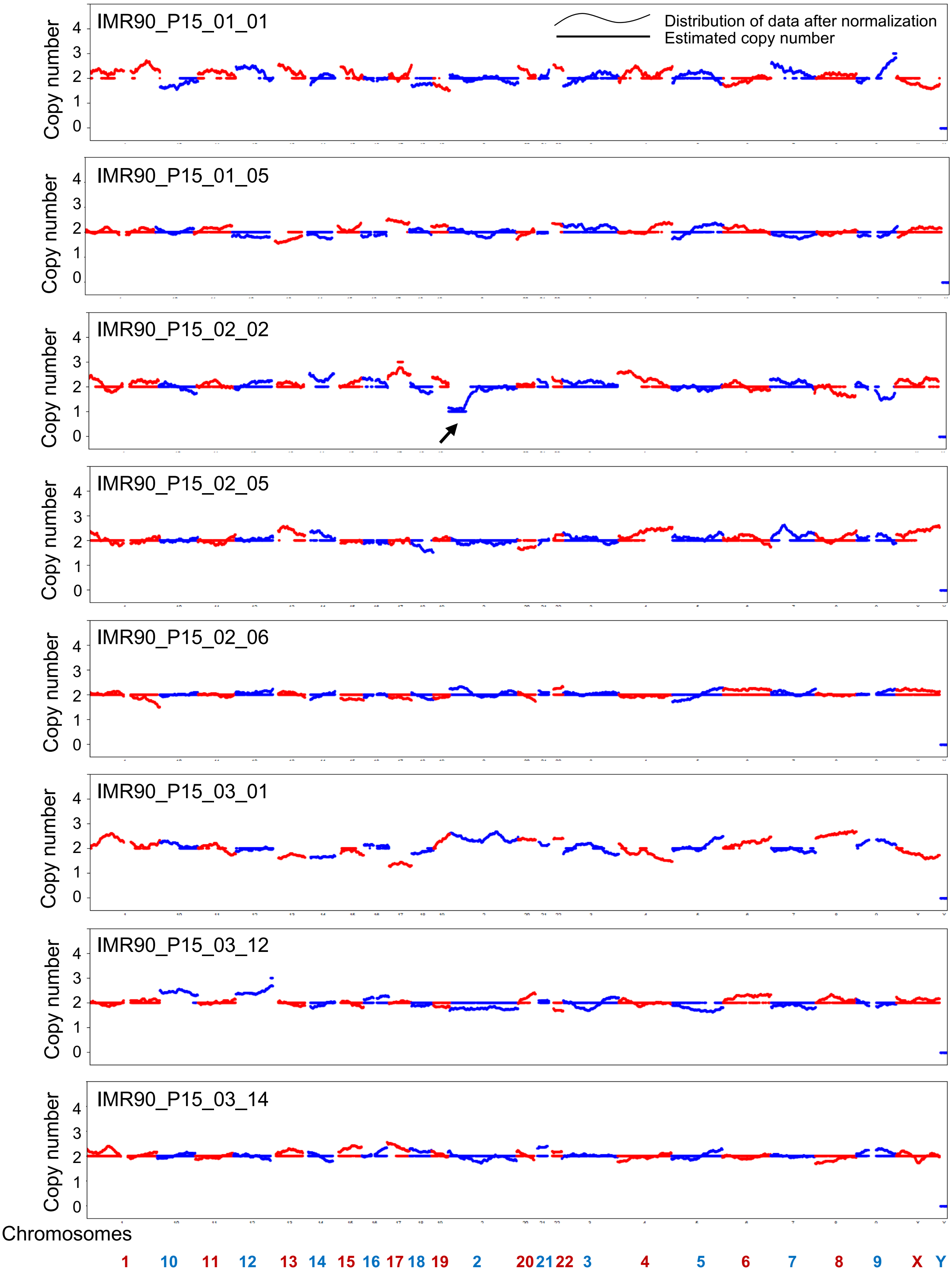


**Supplementary Figure 2.**

Deep senescent cells

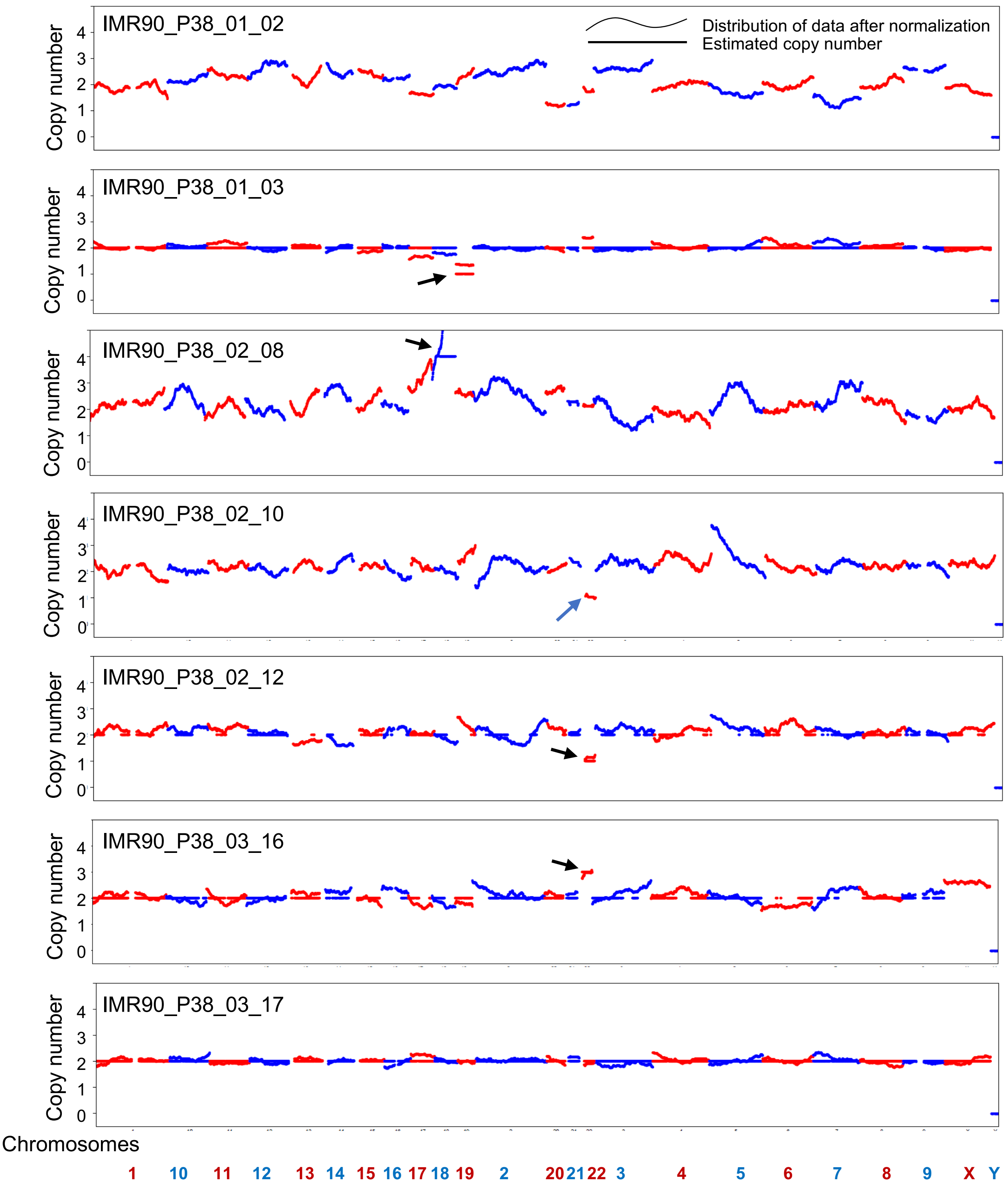


**Supplementary Figure 3.**
